# Disentangling the potential factors defining *Bacillus subtilis* abundance in natural soils

**DOI:** 10.1101/2024.03.11.584434

**Authors:** Xinming Xu, Adele Pioppi, Heiko T. Kiesewalter, Mikael Lenz Strube, Ákos T. Kovács

## Abstract

*Bacillus subtilis* is ubiquitously and broadly distributed in various environments but mostly isolated from soil. Given that species of *B. subtilis* are known as key plant growth-promoting rhizobacteria in agriculture, we here aimed to describe the natural distribution of this species and uncover how biotic and abiotic factors affect its distribution. When comparing different soils, we discovered that *B. subtilis* is most abundant in grasslands, but can rarely be isolated from forest soil, even if the sample sites for the two types of soil are situated in proximity. Differential analysis revealed that spore-forming bacteria exhibited enrichments in the grassland, suggesting niche overlap or synergistic interactions leading to the proliferation of certain *Bacillus* species in grassland environments. Network analysis further revealed that *Bacillus* and other *Bacillota* established a densely interconnected hub module in the grassland soil samples, characterized by positive associations indicating co-occurrence, a pattern not observed in the forest soil. Speculating that this difference was driven by abiotic factors, we next combined amplicon sequencing with physio-chemical analysis of soil samples, and found multiple chemical variables, mainly pH, to affect microbial composition. Our study pinpoints the factors that influence *B. subtilis* abundance in natural soils and, therefore, offers insights for designing *B. subtilis*-based biocontrol products in agricultural settings.

## 1. Introduction

Soil constitutes the primary habitat for numerous members of the *Bacillaceae* family. Its ability to form spores enables survival in adverse conditions and migration to various locations, is believe to be the key factor determining the ecology of these bacteria within the soil ecosystems (Mandic-Mulec et al., 2015). It exerts multifaceted ecological functions in soil. For instance, *Bacillaceae* is capable of breaking down cellulose, hemicellulose and proteins thus helping the degradation of soil organic matter and plant litter (Soares et al., 2012). Moreover, this bacterial family actively participates in nitrogen, carbon, sulfur, and phosphorous cycles within the soil and functions as plant growth-promoting rhizobacteria (PGPR) (Chen et al., 2006; Verbaendert et al., 2011; Mahapatra et al., 2022). Within this family, the species *Bacillus subtilis* is one of the most widely studied PGPR known by its direct modes of action, including nitrogen fixation, phosphorus solubilization, and phytohormones production, among others, and its indirect effects, such as the suppression of soil-borne pathogens (Blake et al., 2021). Several *B. subtilis*-based microbial fungicides or fertilizers have been marketed to improve the plant health and yield of crops (Pérez-García et al., 2011).

The effectiveness and prevalence of PGPR products in field application depend on their adaptation to a spectrum of abiotic and biotic factors (Rilling et al., 2019). Among the abiotic factors, the survival and persistence of endospores formed by *Bacillus spp.* are affected by soil parameters including pH, organic matter content, and calcium levels (Mandic-Mulec et al., 2015). Simultaneously, the dynamic interplay between PGPR and indigenous soil microbiome communities as abiotic factors could potentially determine the success of invasion and colonization by *B. subtilis*. The population density of applied PGPR strain *B. subtilis* B068150 varies among different soil types, with a decrease in the order of clay, loam, and sand (Li et al., 2016). Furthermore, the correlation between the density of *B. subtilis* and with native rhizosphere bacterial communities differs across these soil types. Identifying abiotic and biotic factors that influence the abundance of *B. subtilis* in soil could contribute to the optimization of its effectiveness in plant growth promotion.

In this study, we observed a declining trend of *B. subtilis* colony counts through plate enumeration when we preliminary isolated *B. subtilis* along a transect spanning from a grassland area to a forest. Similarly, a consistently low abundance of *B. subtilis* could be detected in forest soil across five sampled locations in contrast to sample sites in neighboring grassland. To decipher the possible factors contributing to the variation in *B. subtilis* abundance between these two soil types, we investigated the soil physical-chemical properties and the microbial community composition structure. By constructing co-occurrence networks, we compared network characteristics and identified hub taxa in different soil types. We discerned that these hub taxa exhibited altered association patterns through differential network analysis, indicating potentially diverse ecological roles across the soil types. Our work facilitates comprehending the interplay of abiotic and biotic factors that influenced the proliferation of *B. subtilis* in natural environments and gathers insights that might enable the optimization of *B. subtilis*-based PGPR products.

## 2. Material and Methods

### 2.1 Soil sampling and B. subtilis isolation

Soil samples were collected from 5 sample sites distributed across Zealand, Denmark, including Bagsværd (BS) (55.770910 N 12.473349 E), Dyrehaven (DH), Hjortedam Bålplads (HB) (55.774154 N 12.538458 E), Hareskoven (HS) (55.773349 N 12.422606 E), and Frederiksdal Storskov (ST) (55.773444 N 12.422167 E) (Fig. S1B).The primary soil sampling was carried out in Dyrehaven (site DH, Fig S1B), a natural park situated in Zealand, Denmark (55.790829 N 12.566538 E). Here, the sampling was performed along a transect of grassland turning into forest, consisting of six evenly distanced sampling points arranged in line with 30 meters of spacing between each point. Sampling points P6, P11, and P12 were located in the grassland area, while the other three were situated in the forest region (Fig. S1B). Like Dyrehaven, other sampling sites encompass both grassland and forest areas, with the distance between sampling points for obtaining the two soil types less than 500 meters. Samples were collected from a depth of 10 cm below soil surface and placed into sterilized 50 ml tubes. Fresh soil was used for DNA extraction while the remaining soil was stored at 4 °C for subsequent soil property analysis. To isolate *B. subtilis*, 1g of fresh soil sample was suspended in 9 ml sterile distilled water and vortexed for 3 min. To isolate spores from the soil samples, a serial dilution of up to 10,000× was carried out, followed by heat treatment of the soil solution at 80 °C for 10 min. 100 μl of the suspension was spread on lysogeny broth (LB; Lennox, Carl Roth, Germany, Limhamn) medium solidified with 1.5% agar and subsequently incubated at 37 °C for 2 days. Colonies formed a highly structured, rough, opaque, fuzzy white or slightly yellow colony were characterized as *B. subtilis*-like colonies and stored at -80 °C for further phylogeny determination. Full-length of 16S rRNA gene was PCR-amplified with primer pair 27F (5’-AGAGTTTGATCCTGGCTCAG-3’) and 1492R (5’-GGTTACCTTGTTACGACTT-3’) and sent out for sanger sequencing. The 16S rRNA sequences of each strain were then taxonomically assigned using NCBI-blastn (Fig. S1A).

### 2.2 Soil chemistry

Soil pH was measured using a pH meter with a soil-to-water ratio of 1: 2.5. Soil moisture content was calculated gravimetrically on a wet weight basis after oven-drying at 60 °C for 48 h. Available magnesium (Mg) and potassium (K) ions were extracted using ammonium acetate solution and quantified through flame photometrically. Soil phosphorus (P) content was extracted using sodium bicarbonate solution and measured by spectrophotometry. Soil particle size distribution was assessed and particles < 0.002mm, 0.002-0.02 mm, 0.02-0.2 mm, and 0.2-2.0 mm were classified as clay, silt, fine sand, and coarse sand, respectively. Total soil carbon (C) and nitrogen (N) were determined through dry combustion using a CN analyzer. The soil properties analysis were performed by AGROLAB GmBH (Landshut, Germany).

### 2.3 DNA extraction and amplicon sequencing

Genomic DNA was extracted from 250 mg soil using DNeasy PowerSoil kit (Qiagen, Hilden, Germany) according to the manufacturer’s instructions. The concentration and quality of DNA were evaluated by NanoDrop DS-11+ Spectrophotometer (DeNovix, Wilmington, U.S.) and Qubit 2.0 Fluorometer (Thermo Fisher Scientific, Massachusetts, U.S.). From each sampling site, gDNA was extracted in in three biological replicates. To assess the bacterial and fungal communities, the V3-V4 hypervariable region of bacterial 16S rRNA gene and ITS2 region of fungi were amplified using the primer pairs of 341F (5’-CCTACGGGNGGCWGCAG-3’) and 805R (5’-GACTACHVGGGTATCTAATCC-3’), and gITS7ngs (5’-GTGARTCATCRARTYTTTG-3’) and ITS4ngs (5’-TCCTSCGCTTATTGATATGC-3’) (Frank et al., 2008; Tedersoo and Lindahl, 2016), respectively. Thermal cycling conditions for 16S rRNA amplification were as follows: 95 °C for 15 min followed by 30 cycles of 95 °C for 30 s, 62 °C for 30s and 72 °C for 30s and a final extension at 72 °C for 5min. For ITS amplification, PCR program included initial denaturation for 15 min at 95 °C; 30 cycles of 45 s at 95 °C, 45 s at 50 °C and 90 s at 72 °C; and a final extension for 5 min at 72 °C. 25 μl PCR reactions contain 10.6 μl DNase-free water, 12.5 μl TEMPase Hot Start 2× Master Mix, 0.8 μl of each primer (10 μmol/L), and 0.3 μl of DNA template. The PCR products were purified using NucleoSpin Gel and PCR Cleanup Kit (Macherey-Nagel), following the manufacturer’s instructions. After purification, the products were pooled at equimolar concentrations. The bacterial and fungal amplicon pools were submitted to Novogene Europe Company Limited (United Kingdom) for sequencing on a NovaSeq PE250 platform with 2 million reads per sample. The raw sequence data has been deposited to the NCBI Sequence Read Archive (SRA) database under the BioProject accession number PRJNA1012666.

### 2.4 Sequencing data processing analysis

Raw sequence data was processed with the QIIME2 pipeline (Bolyen et al., 2019). Primers and barcodes were removed with the QIIME2 plugin cutadapt and after demultiplexing, filtered reads were denoised, merged and chimera-checked using DADA2 (Callahan et al., 2016). In total, we retained 1,121,356 reads and 1,330,258 reads for 16S rRNA and ITS with an average of 37,774 and 44,341 reads for each sample, respectively. Taxonomy was assigned to sequence using the Naive Bayesian classifier trained on the SILVA (132 release) reference database for 16S rRNA and the UNITE reference database for ITS, respectively (Quast et al., 2013; Nilsson et al., 2019). All the ASVs annotated as members of the *Bacillus* genus were then further aligned by use of BLASTN to a custom *B. subtilis* V3-V4 database constructed by RibDif (Strube, 2021). ASVs with similarities at >95% were assigned as *B. subtilis*.

### 2.5 Statistical analyses

All statistical analyses were conducted using R (version 4.1.1). The number of sequences in each sample was rarefied corresponding to the sample with the fewest reads, resulting in 6961 reads for 16S rRNA and 7999 reads for ITS analysis per sample (Fig. S2). To determine genera that exhibit differential abundance between soil types, we employed the analysis of composition of microbiomes with bias correction (ANCOM-BC), which considers the compositional nature of microbiome sequencing data. To implement this method, the ANCOM-BC v1.4.0 R package was applied to raw read count tables, as it incorporates internal correction for library size and sampling biases (Lin and Peddada, 2020). The significance cutoff was set at an adjusted p-value of 0.05, p values were adjusted using the Bonferroni method. The cumulative count of differentially abundant genera found in soil environments was summarized in a balloonplot using the ggballoonplot function from the ggpubr R package (Kassambara, 2023). For multivariate analysis, Bray-Curtis index was chosen as a dissimilarity measure between ASVs and were visualized through non-metric multidimensional scaling (NMDS) ordination plots using the ‘metaMDS’ function from the ‘vegan’ package (Dixon, 2003). Furthermore, a permutation test was performed using the ‘envfit’ function to fit environmental metadata vectors to the microbial community composition (Dixon, 2003). The correlation of each environmental variable with bacterial diversity, fungal diversity, and *B. subtilis* ASVs were assessed by the Mantel test using the ‘vegan’ package with 9,999 permutations (Dixon, 2003). The pairwise correlation among different environmental variable was determined by Pearsons correlations and visualized with the linkET package in R (Yun, 2023). For univariate analyses, students t-tests were performed to identify soil physical-chemical properties that exhibited significant differences in specific soil types. Shannon alpha diversity was determined using the R package ‘ampvis2’ based on the rarefied data (Andersen et al., 2018).

### 2.6 Construction of co-occurrence networks

To infer keystone taxa in different soil types, co-occurrence networks were constructed using the ‘NetComi’ package (Peschel et al., 2021). The SparCC method was employed for network construction, which is specifically designed for compositional data (Weiss et al., 2016). Genera with read counts exceeding 100 in at least 5 samples while adhering to a sparsity threshold of 0.6 were used to generate the correlation matrix. The netAnalyze function was employed for network characterization to identify the modularity and clusters within the network. Hub taxa were defined based on the eigenvector centrality value exceeding the empirical 95% quantiles. These hub taxa termed as “keystone taxa”, were characterized by high degree and closeness, together with low betweenness centrality, which were previsouly proposed as keystone taxa indicators. Quantitative comparisons between networks were executed using the ‘netCompare’ function implemented in ‘NetCoMi’, involving a permutation-based approach with an adaptive Benjamini-Hochberg correction for adjusting p-values to account for multiple testing. A differential association network was constructed using the ‘diffnet’ function where two nodes are connected if they are differentially associated between two environments.

## 3. Results

### 3.1 Bacterial and fungal community composition in grassland and forest soil

Members of the *Bacillaceae* family was isolated from soil samples after heat treatment that selects for spore formers. In particular, members of the *B. subtilis* species complex was easily identified on nutrient-rich cultivation media as this species forms highly structured colonies. We observed that colonies of spore formers could be isolated corresponding to *B. subtilis* or closely related species from the grassland soil and hardly detectable in the samples originating from the forest (none *B. subtilis* colonies at sampling site P15), even if the sample site was just few hundred meters from apart (Table S2).

We then aimed to obtain a comprehensive overview of the microbial community composition across all five sampling sites and assess the distinctiveness of *B. subtilis* enrichment in the grassland area. We found the predominant bacterial phyla observed across the sampling sites were *Pseudomonadota* (*Proteobacteria*), *Actinomycetota* (*Actinobacteria*), *Planctomycetota* (*Planctomycetes*), and *Bacillota* (*Firmicutes*). The abundance of *Bacillota* in grassland was higher than that in forest soil except for sampling site BS (Fig. S3A). *Ascomycota* and *Basidiomycota* stood out as the dominant fungi across all sampling sites, collectively accounting for over 50% of the fungal community (Fig. S4B). At the family level microbial composition, *Xanthobacteraceae*, *Chthoniobacteraceae*, and *Isosphaeraceae* were prevalent in most samples (Fig. 1A). The grassland at sampling site Dyrehaven (DH_G) exhibited the highest abundance (11.1%) of *Bacillaceae* among all the samples. While most sampling sites demonstrated a higher *Bacillaceae* abundance in grassland compared with the forest soil, BS_F displayed a lower abundance of 2.3% in grassland compared to 0.9% in forest soil. Notably, *Aspergillaceae* constituted the most abundant fungi in sample DH_G (19.6%) (Fig. 1B). *Russulaceae* were barely detected in the grassland but displayed an abundance of 31.3% in the forest soil at Hareskoven and 10.2% at Dyrehaven. *Mortierellaceae*, *Trimorphomycetaceae*, and *Sporomiaceae* were widespread in most samples, contributing to the overall fungal diversity.

**Figure 1.**
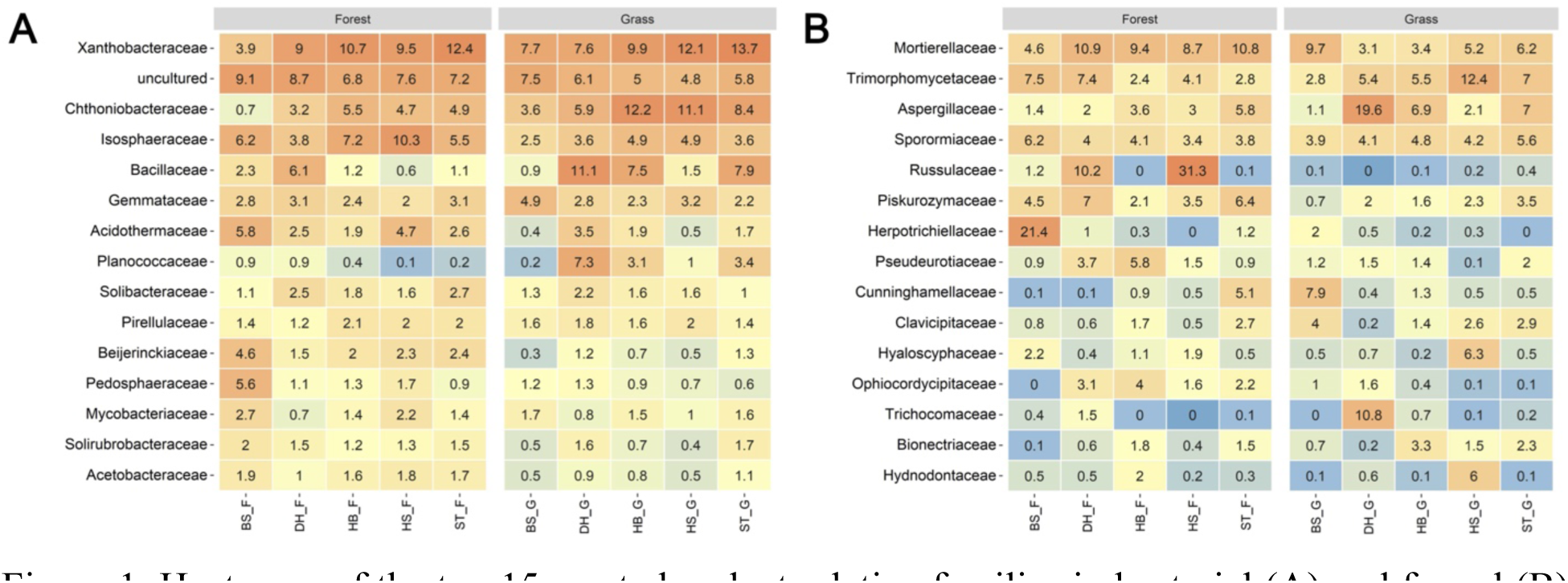
Heatmaps of the top 15 most abundant relative families in bacterial (A) and fungal (B) communities, respectively. The left panel showcases samples collected from forest soil, while the right panel displays samples from grassland.

### 3.2 Unveiling specific taxon enrichment in grassland through differential analysis

To comprehensively assess taxon presented similar enrichment patterns in grassland as *B. subtilis*, we employed the ANCOM-BC model to analyze the differential relative abundance between the two soil types (Lin and Peddada, 2020). Our analysis revealed a substantial increase in the relative abundance of specific bacterial taxa within the grassland environment (Fig. 2A). While annotating bacterial taxa enriched across various samples on the phylogenetic tree, we identified that families such as *Bacillaceae* and *Hyphomicrobiaceae* displayed a consistent pattern of enrichment across four out of five samples (Fig. 2B). Delving into the specifics, genera such as *Streptomyces*, *Candidatus Udaeobacter*, and *Rhodanobacter* demonstrated a notable increase in their relative abundance within the grassland habitat. Furthermore, when surveying the differential abundance within the *Bacillaceae* family, we found that several genera, *Ammoniphilus*, *Bacillus*, *Chungangia*, *Lysinbacillus*, *Psychrobacillus*, and *Sporosarcina* exhibited significant enrichment across multiple samples, except for the BS sampling site (Fig. 4C).

**Figure 2.**
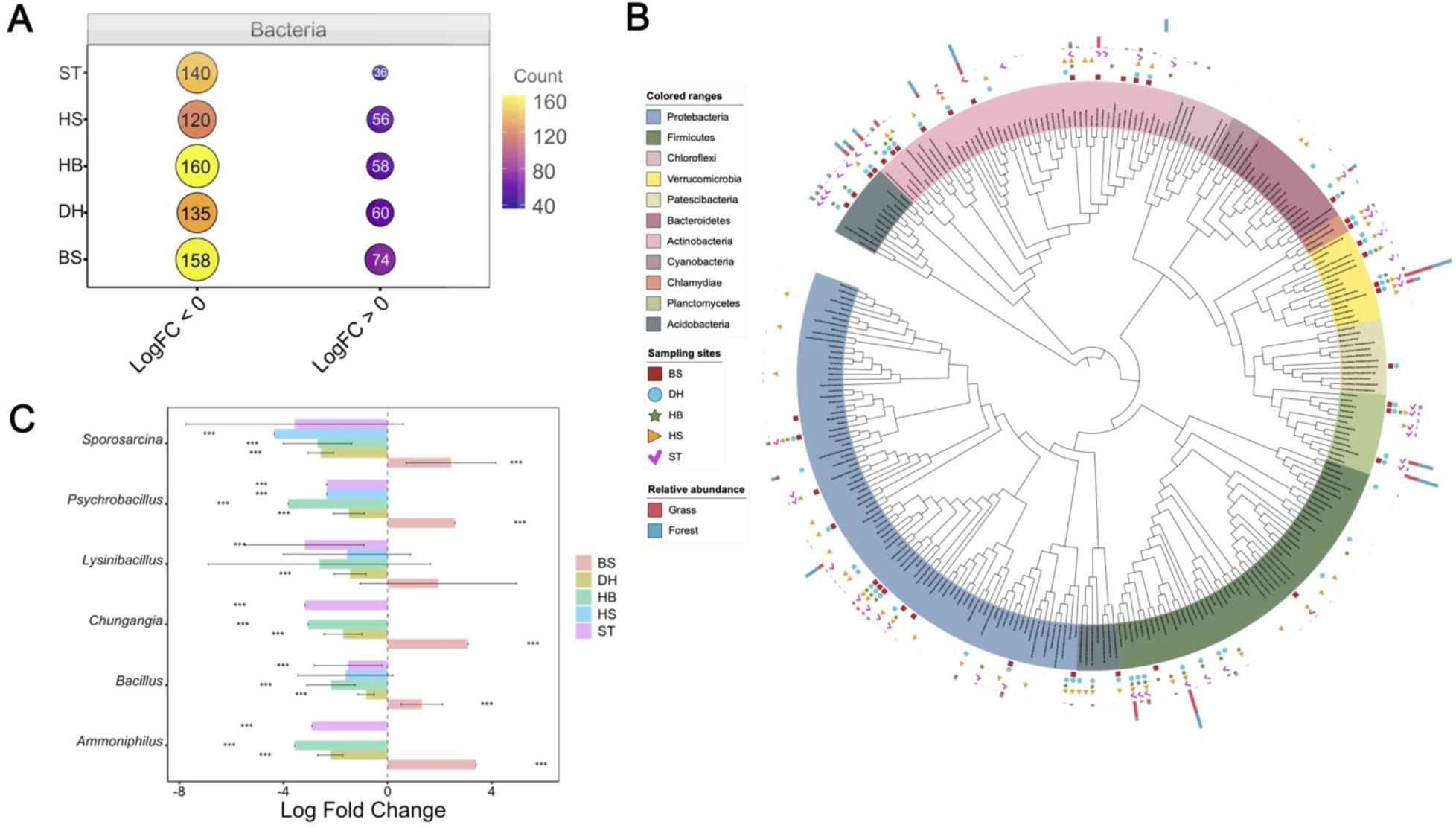
(A) Bubbles representing the size equivalent to the accumulated sum of genera that are enriched in respective soil types. LogFC < 0 indicates genera enriched in grassland, LogFC >0 indicates genera enriched in forest soil. (B) Phylogenetic tree of all ASVs detected across samples. The rings, from the inside to the outside, represent taxa enriched in sampling sites BS, DH, HB, HS, and ST (square, circle, star, triangle, and check mark, respectively). The outer ring indicates the average cumulative normalized abundance of each ASV in grassland (red) versus forest soil (blue). (C) Differential abundance analyses stratified by soil type using ANCOM-BC. Data are represented by effect size (log fold change) and 95% confidence interval bars (two-sided; Bonferroni adjusted) derived from the ANCOM-BC model. All effect sizes with adjusted p < 0.05 are indicated, *significant at 5% level of significance; **significant at 1% level of significance; ***significant at 0.1% level of significance.

### 3.3 Co-occurrence patterns of bacterial and fungal soil communities under different soil types

Having gained a comprehensive overview of the microbial community composition and identified differential taxa between the two soil types, we employed network analysis to further investigate the co-occurrence patterns within the soil microbial communities across distinct soil types. The topological characteristics of the network including modularity, clustering coefficient, and natural connectivity among bacterial taxa, exhibited higher values within the grassland than the forest soil. The bacterial community in grassland forms a network with lower density but a higher proportion of positive edges, accounting for 61.39 %, in contrast to the forest soil network, which exhibits a positive edge proportion of 55.06 %. Calculations of centrality measures including betweenness, degree, closeness, and eigenvector centrality identified hub taxa in the grassland predominantly belong to the *Bacillota* phylum and encompass genera such as *Ammoniphilus*, *Bacillus*, *Chungangia*, and *Sporosarcina.* Together with the genus *Acidothermus*, these taxa formed a densely interconnected module. Interestingly, the assembly of spore-forming taxon did not exhibit the same prominence in the forest soil network. Differential network analysis revealed that the connectivity pattern of these spore-forming bacteria transformed from positive connections to negative connections in the forest-derived samples (Fig. 3). Hub taxa in the forest soil included members of the verrucomicrobial clade *Candidatus Udaeobacter* and *Candidatus Xiphinematobacter*, as well as bacteria belonging to *Methyloligellaceae, Planctomycetale*s, and genus *Bauldia*.

**Figure 3.**
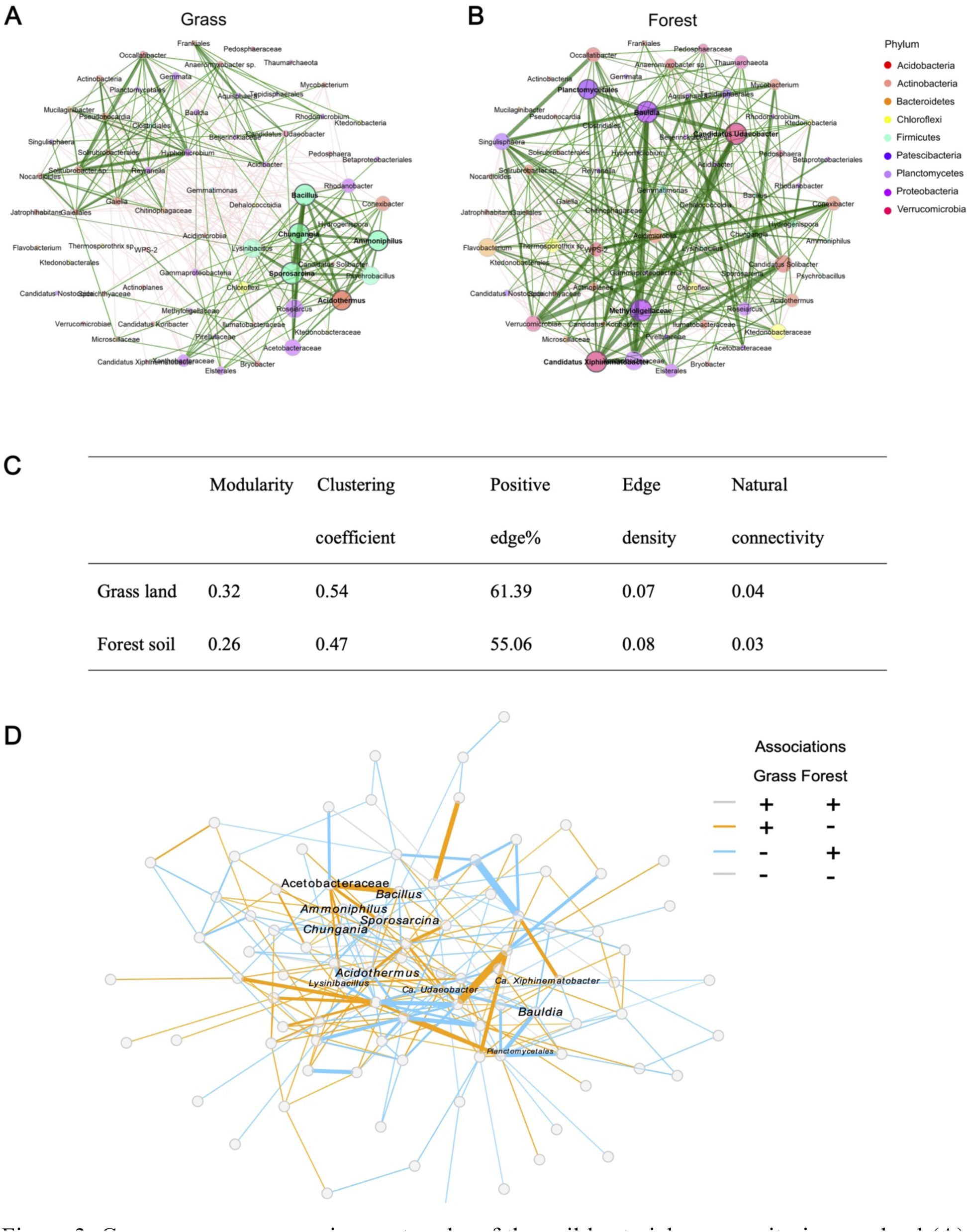
Co-occurrence comparison networks of the soil bacterial community in grassland (A), and forest soil (B). Nodes represent ASVs aggregated to the genus level and are colored according to the respective phylum. ASVs that were not assigned on the genus level were annotated at a higher taxonomy level. Edges in the networks represent positive (green) and negative (red) correlations > |0.6| calculated by the SparCC method. Edges thickness is proportional to partial correlation. Eigenvector centrality of each node was used to define hub taxa and highlighted in bold. (C) Topological characteristics of global network. (D) Bacterial differential network. Nodes are connected if they exhibit differential associations between two groups. Orange lines indicate taxa that were positively associated in grassland but negatively associated in the forest. Blue lines represent taxa that were negatively correlated in the grassland but positively in the forest. Grey lines indicate taxa that maintained consistent correlations across both environments.

Within the fungal network, hub taxa comprised dominant genera present across the samples. Specifically, in the grassland network, *Solicoccozyma*, *Sporormiella*, and *Trimmatostroma* were identified as hub taxa, whereas *Meliniomyces*, *Pochonia*, and *Russula* were highlighted as ‘keystone’ taxa in the forest network (Fig. S4A&B). *Sporormiella* exhibited a substantial high degree and closeness in grassland, presenting as a ‘central hub’ taxon and closely connected with other hub taxa. Furthermore, the forest fungal community network contained more distinct modules, a slightly higher proportion of positive connections (57.14 %) between taxa, and a higher clustering coefficient. These findings collectively provide insights into the intricate dynamics of microbial interactions under different environments.

### 3.4 Correlation between bacterial and fungal community and soil physiological properties

To identify the biotic and abiotic factors that influenced the differentially distributed microbiome in grassland and forest, we analyzed the correlation between bacterial and fungal communities with soil physiological properties. We first inspected the bacterial and fungal diversity in the soil samples originating from grassland and forest sites. The Shannon index of bacterial richness was lower in the grassland than in the samples from the forest soil. In contrast, fungal abundance exhibited an opposite trend (Fig. 4A). Nonetheless, these variations do not adhere a general trend and are contingent upon sampling sites (Fig. S5). Prior to analyzing the physicochemical correlations with the microbiome, we conducted a preliminary analysis for correlations. A few physicochemical parameters were excluded as strong positive correlations among organic carbon, soil nitrogen content, and organic substance since they would potentially overinflate the effect on the communities. Mantel tests revealed that bacterial diversity was highly related to soil pH while multiple factors, including pH, soil magnesium, and soil texture significantly impacted the fungal richness (Fig. 4B). Furthermore, notable correlations were observed between environmental variables such as pH, soil calcium, and phosphorus content with *B. subtilis*-assigned ASVs.

**Figure 4.**
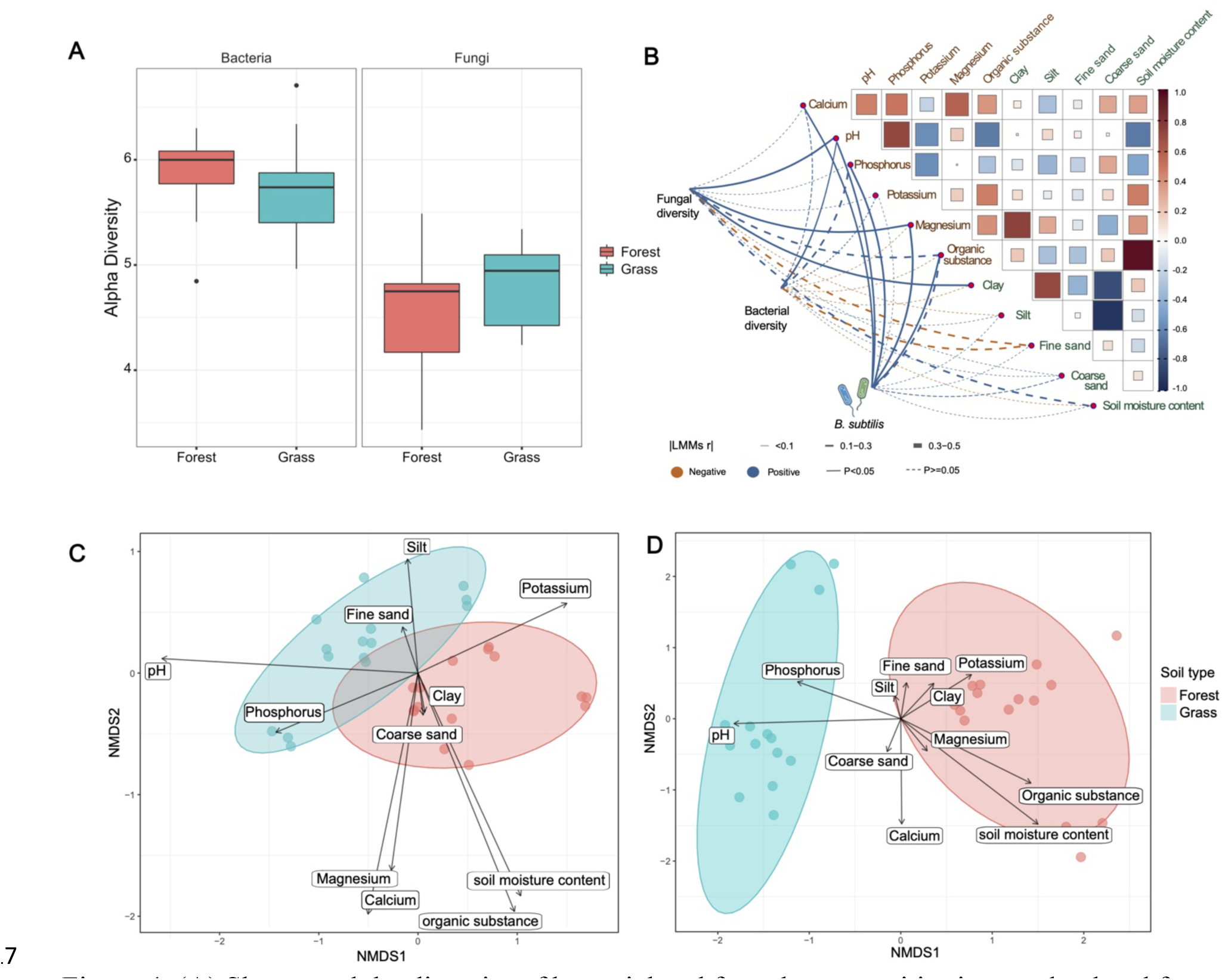
(A) Shannon alpha diversity of bacterial and fungal communities in grassland and forest soil. (B) Mantel test analyzed the relationships between soil properties, microbial diversity and *B. subtilis* ASVs. The thickness of lines is proportional to correlation, blue lines indicate positive correlations, orange lines indicate negative correlations. Solid lines represent significant correlations, dash lines represent non-significant correlations. Pearson’s correlations were calculated between soil physicochemical properties. Beta diversity of bacterial (C) and fungal (D) communities were calculated with the Bray-Curtis dissimilarity and visualized as circles in NMDS (bacterial community: stress = 0.116, adonis: R^2^ = 0.153, p = 0.001; fungal community: stress = 0.177, adonis: R^2^ = 0.146, p = 0.001). Vectors represent soil physicochemical properties with lengths proportional to the level of correlation.

The NMDS visualization of microbial communities using Bray-Curtis distances effectively depicted separations originating from different soil types and statistically verified by adonis (bacterial community: stress = 0.116, adonis: R^2^ = 0.153, p = 0.001; fungal community: stress = 0.177, adonis: R^2^ = 0.146, p = 0.001). Chemical and physical indicators of soil properties correlated with the community structure are shown as vectors on the NMDS ordinations (Fig. 4C&D). Specifically, soil pH, organic substance and moisture content were related to the differences in microbial community structure between soil types. Furthermore, the soil potassium content significantly correlated with the distinct bacterial community structure. Among these factors, soil pH exerted the most pronounced influence on shaping unique patterns within the bacterial community while soil moisture content affected the fungal community primarily.

## 4. Discussion

Despite numerous studies that have thoroughly examined the abiotic factors that would promote the growth of *B. subtilis* in the laboratory, all reported experiments were conducted under well-controlled conditions, leaving a huge gap between *in vitro* laboratory conditions and field applications. In this study, we observed a natural gradient feature varying degrees of *B. subtilis* abundance across grassland and forest soil. We then isolated soil samples from locations encompassing both grassland and forest soil at close distances and investigated the soil properties and microbiome communities. Our results extend the knowledge about the factors that influence *B. subtilis* survival and abundance *in situ* and the mechanisms shaping bacterial and fungal soil community structure, potentially enhancing the efficacy of *B. subtilis* application in agricultural scenarios.

### 4.1 Spore-forming bacteria were enriched in grassland

The amplicon data were subjected to more in-depth analysis to reveal whether *B. subtilis* was the sole microbiome components that was enriched in the grassland, or if multiple taxa exhibit the same presence-absence pattern across both environments. Together with co-occurrence network, we found that hub taxa detected in the grassland, particularly spore-forming bacteria, exhibit substantially higher abundance in grassland compared to the forest. The formation of endospores is affected by several factors, including environmental pH, and the presence of carbon, nitrogen, and phosphorus sources (Mandic-Mulec and Prosser, 2011). The generally lower mineral and water content in grassland may contribute to the enrichment of spore-forming bacteria (Logan and De Vos, 2011). This enrichment could potentially serve as a mechanism to adapt to the harsher environments present in the grassland.

To our surprise, the abundance of *B. subtilis* is not associated with the overall abundance of the *Bacillus* genus. In most sampled sites, the *Bacillus* genus exhibited significantly higher abundance in the grassland than in forest soil, except for sampling site Bagsværd. One possible reason is the abnormally high soil water content, with an average value of 25.98 %, reaching as high as 63.93 % at this specific site, and an average organic content of 4.85 % but escalated to 14% at Bagsværd. Future experiments on removing spore-forming bacteria or manipulating abiotic factors will provide further insights into its influence on the abundance of *B. subtilis*.

### 4.2 Co-occurrence network indicates that the microbiome exhibits different association patterns between two soil types

Microbial co-occurrence networks have become a standard tool in the analysis of microbiome composition and to infer significant associations between pairs of co-occurring taxa and ascribe to biological interactions, such as co-existence or mutual exclusion (Faust and Raes, 2012; Fuhrman et al., 2015; Pérez-Valera et al., 2017). Since microbial interactions are likely to change between soil types, we integrated NetComi to construct co-occurrence networks that allow us to compare the network structures and the differential networks and to identify taxon differentially associated between the grassland and forest soil groups (Peschel et al., 2021).

The clustering coefficient in a network quantifies the degree to which nodes in a local neighborhood tend to be interconnected, where stronger microbial interaction and more dynamic bacterial communities were observed in grassland. Grassland has high modularity indicating the network has dense connections within certain taxon groups and sparse connections between these groups, reflecting possible niche differentiation or environmental heterogeneity. The forest bacterial network exhibits a slightly higher density compared to grassland, accompanied by a greater number of negative connections that might arise from differential niche adaptation and/or competition (Faust and Raes, 2012; Faust et al., 2012; Dohi and Mougi, 2018).

Previous studies have highlighted that soil abiotic factors are the main drivers of bacterial community structure, composition, and topological features of co-occurrence networks across biogeographic scales (Martiny et al., 2006; Agler et al., 2016; Fierer, 2017; Meyer et al., 2018; Goberna et al., 2019). Goberna et al. specifically identified soil parameters like oxidizable carbon, total nitrogen, moisture, and pH as essential abiotic factors in determining the spatial distribution of soil bacteria and their co-occurrence patterns (Goberna et al., 2019). In our study, the formation of the hub module primarily composed of spore-forming bacteria displayed substantial connectivity within the grassland soil, a phenomenon not observed in the forest soil. This implies a transition in the ecological roles of these bacteria across the two distinct soil types, suggesting an alteration within bacterial interactions and potential functions in respective ecosystems. The genera *Chungangia*, *Sporosarcina*, and *Ammoniphilus* have lower than 1% relative abundance in grassland but together with *Bacillus,* formed a densely interconnected hub module characterized by positive associations. In a co-occurrence network, positive interactions could result from cross-feeding, co-colonization, or niche overlap. The positive association of the *Bacillus* genus and these low-abundant spore-forming bacteria in the bacterial community could be due to niche overlap where they share similar environmental preferences (soil pH, moisture content, organic matter). Simultaneously, a positive correlation may indicate synergistic interactions where the metabolites secreted by one taxon are consumed by another (Freilich et al., 2011; Berry and Widder, 2014; Kodera et al., 2022). Thus, the hub taxa in grassland may benefit from the biochemical activities of others, leading to the proliferation (including *B. subtilis*) in the grassland environment.

Members of the verrucomicrobial clade *Ca. Udaeobacter* are among the most dominant bacterial phylotypes in soil. The provisional designation “*Candidatus*” refers to bacteria that have not yet been successfully cultured and grown in the laboratory, requiring a more complete description (Murray and Stackebrandt, 1995). Previously, a study investigated the association between the abundance of *Ca. Udaeobacter* and soil pH by sampling 150 forest and 150 grassland soils (Willms et al., 2021). The reported data is consistent with our finding that *Ca. Udaeobacter* has the highest abundance in strongly acidic soil, particularly within forested regions. Furthermore, *Ca. Udaeobacter* together with *Ca. Xiphinematobacter*, *Planctomycetales*, *Methyloligellaceae*, and *Bauldia* were quantified as hub taxa in the forest bacterial network. However, our current understanding of these bacterial taxa remains limited, leaving only preliminary insights into their associations within the forest ecosystem.

Generally, fungal communities are tightly correlated with plant diversity and composition (Yang et al., 2017). Meanwhile, biotic factors were suggested to play more important roles than abiotic factors for fungal network assembly compared to bacterial network (Yang et al., 2017; Adamczyk et al., 2019). The clustering coefficient and positive edge proportion of the fungal network were similar between the two soil systems. The fungal forest network displayed a higher modularity suggesting that groups of fungi share specialized ecological niches or function roles. Nevertheless, fungal-bacterial relationships remain to be explored. Future progress in understanding individual fungal-bacterial interactions might help us to understand the intricate networks in grasslands and forests.

Although co-occurrence network construction has been widely applied and allows the extraction of simple patterns from complex datasets, follow-up analysis on the networks requires integration of laboratory experiments that go beyond network analysis and characteristic interpretation (Ling et al., 2016; Marta Goberna and Miguel Verdú, 2022). To validate the regulatory effect of hub taxa, experiments need to be conducted involving the removal of keystone microbial species or modification of interactions between microbiomes that allow a better understanding of the microbiome functions and manipulation of soil microbiome communities.

### 4.3 Multiple abiotic factors shape the microbial community structures in soil

The richness of the soil microbiome exhibits substantial variation between the grassland and forest soil types. However, no consistent trend indicates that one soil type has a higher richness than the other. Contrary to our findings, previous studies have suggested that grasslands contain richer bacterial diversity than forests (Delgado-Baquerizo and Eldridge, 2019). Yet, little is known about various environmental factors and the spatial scale heterogeneity that can influence soil microbial diversity (Vos et al., 2013). Based on the Mantel test, we identified that both bacterial and fungal richness were highly related to soil pH. A comparative analysis of soil samples collected within the Earth Microbiome Project suggested that environmental factors such as pH might outweigh the plant biomes in shaping soil alpha diversity, implying a diverse array of abiotic parameters could contribute to the observed variations in alpha diversity in our results (Rousk et al., 2010; Wardle and Lindahl, 2014; Walters and Martiny, 2020).

Despite the relatively short distance between the grassland and forest areas, we observed a strong effect of abiotic factors on microbial community structure. The divergence in beta diversity of fungal community structure and composition between soil types exceeded that of the bacterial community, which may be attributed to the closer interaction of fungi with plants, given their roles as root symbionts, endophytes and pathogens (Peay et al., 2013; Vos et al., 2013). Both soil pH and moisture content emerged as pivotal factors in shaping the composition of both bacterial and fungal communities. Meanwhile, we found that soil pH significantly affected bacterial community composition while exerting a lesser influence on the fungal community. This pattern is consistent with the idea that bacteria thrive in narrow pH ranges, whereas fungi tend to tolerate wider pH ranges for growth (Lauber et al., 2009; Rousk et al., 2010; Griffiths et al., 2011; Kaiser et al., 2016). Rousk and colleagues argued that the observed correlation between soil pH and fungal community composition is mediated by the dynamics of the bacterial community and its competitiveness along the pH gradient (Rousk et al., 2010). Therefore, soil pH may serve as one of the main drivers, together with other environmental factors, in shaping the community changes between grassland and forest directly or indirectly.

Li et al. observed low population densities of the inoculant *B. subtilis* B068150 in sandy soil and attributed this to low levels of organic matter and reduced water potential (Li et al., 2016). Similarly, we have identified significant correlations between *B. subtilis*-assigned ASVs and soil organic substances, soil pH, as well as soil minerals such as phosphorus, magnesium, and calcium. Numerous laboratory tests have effectively demonstrated the interplay between mineral contents and the growth of *B. subtilis*, for instance, the effect of magnesium on its modulation of cell division frequency and calcium contents on biofilm dispersion (Mhatre et al., 2017; Nishikawa and Kobayashi, 2021; Guo and Herman, 2023). Meanwhile, as a PGPR strain, the effect of *B. subtilis* on solubilization and enhancement of magnesium and calcium content in both soil and plants is known (B. O. Sun et al., 2020; B. Sun et al., 2020; Mahapatra et al., 2022). Nonetheless, research is still limited on whether soil abiotic properties modulate the performance of *B. subtilis*.

A few studies have tapped into the soil properties that influence other species of the *Bacillus* genus. Ramírez et al. demonstrated that soil phytate availability determines the performance of PGPR inoculant *Bacillus amyloliquefaciens* FZB45, revealing that FZB45 only promoted plant growth and phosphorus uptake at a high rate of soil phytate (Ramírez and Kloepper, 2010). The diversity of soil indigenous communities is one of the biotic factors that influence the survival of inoculated *Bacilli*. Mallon et al. proposed the principle of the diversity-invasibility relationship, wherein PGPR inoculation, as a form of human-induced microbial invasion, is less likely to succeed in highly diverse communities when available resources and niches for invaders are limited (Mallon et al., 2015). This theory has been tested on the *Bacillus* genus (*Bacillus pumilus* and *Bacillus mycoides*) to assess its applicability but discovered the principle does not seem to comply throughout all spore-forming bacteria (Mawarda et al., 2022). Thus, there is no clear answer to which abiotic or biotic factors might primarily determine the proliferation of the members of the *Bacillus* genus in soil. In this study, we leveraged a natural gradient feature varying degrees of *B. subtilis* abundance to elucidate the biotic and abiotic factors influencing its presence. Future experiments on modulating these factors independently and collaboratively will aid in predicting the outcomes of PGPR application and enhancing its efficacy.

## 5. Conclusion

In this study, we observed a pattern where *B. subtilis* has a lower abundance in forest soil than in grassland despite their close proximity. Our results highlight distinct patterns in the microbiome community structure between two soil types, with significant influence from abiotic factors such as soil pH, moisture content, and organic substances. Co-occurrence network analysis demonstrated differing network topology characteristics between the two soil types. In grassland, *Bacillus*, *Ammoniphilus*, *Chungangia*, and *Sporosarcina,* and *Acidothermus* formed a positively interconnected hub module, a pattern that was absent in the forests. Furthermore, differential analyses revealed a notably higher abundance of spore-forming bacteria within the hub module in the grassland, suggesting a potential niche overlap or cross-feeding among these bacteria.

## Declaration of competing interest

The authors declare that they have no known competing financial interests or personal relationships that could have appeared to influence the work reported in this paper.

## Acknowledgments

X.X. was supported by a China Scholarship Council fellowship. This project was supported by the Danish National Research Foundation (DNRF137) for the Center for Microbial Secondary Metabolites and the Novo Nordisk Foundation within the INTERACT project of the Collaborative Crop Resiliency Program (NNF19SA0059360).

## Notes

### Competing Interest Statement

The authors have declared no competing interest.

