## Supplementary material for "Disentangling the potential factors defining *Bacillus subtilis* abundance in natural soils": Table S1-S2 and Fig S1-S5

**Supplementary tables**

Table S1. Soil physicochemical variables aggregated across all disturbed and reference soil samples. The mean values were compared using t-tests and shown as means ± standard deviation. Asterisks indicate significantly higher values at the following significance levels: * $P < 0.05$, ** $P < 0.01$, *** $P < 0.001$.

| **Variable** | **Grassland** | **Forest** |
|---|---|---|
| pH | 5.28 ± 0.30 *** | 4.66 ± 0.41 |
| Calcium (mg/kg) | 1287.40 ± 532.58 | 1434.20 ± 178.07 |
| Phosphorus (mg/100g) | 2.76 ± 1.83** | 1.40 ± 0.46 |
| Potassium (mg/100g) | 8.32 ± 2.19 | 10.6 ± 1.65 |
| Magnesium (mg/100g) | 4.92 ± 1.05 | 7.18 ± 3.16 |
| Organic substance (%) | 3.98 ± 0.7 | 7.36 ± 4.67 ** |
| Clay (%) | 9.12 ± 1.44 | 10.1 ± 1.75 |
| Silt (%) | 9.54 ± 2.14 | 10.32 ± 1.43 |
| Fine sand (%) | 33.56 ± 8.32 | 72.68 ± 72.07 |
| Coarse sand (%) | 43.64 ± 14.15 | 40.48 ± 14.8 |
| Organic Carbon (%) | 2.30 ± 0.15 | 4.98 ± 1.39 |
| Total Nitrogen (%) | 0.196 ± 0.1 | 0.29 ± 0.15 |
| Soil moisture content (%) | 20.27 ± 4.31 | 38.15 ± 17.98*** |

Table S2. *B. subtilis* colony counts in each sampling sites determined by 16S rRNA sequencing.

| | BS | DH | HB | HS | ST |
|---|---|---|---|---|---|
| Grassland | 3 | 3 | 3 | 4 | 6 |
| Forest | 1 | 0 | 0 | 0 | 0 |

### Supplementary figures

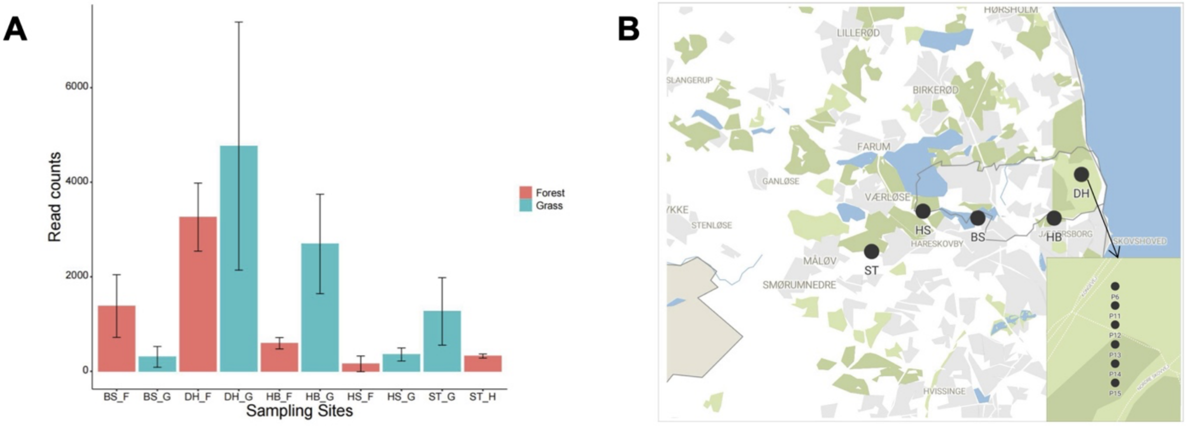


**Fig. S1** (A) Read counts of *B. subtilis* ASVs in each sampling sites. (B) Locations of each sampling sites.

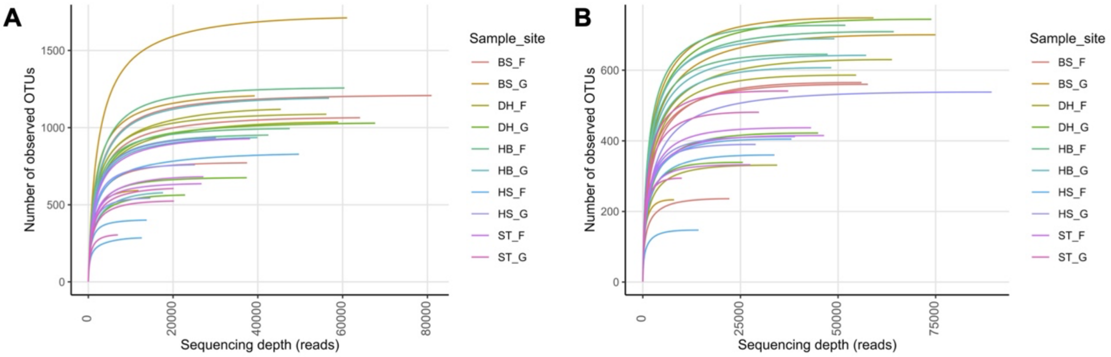


**Fig. S2** (A) Bacterial and (B) Fungal rarefaction curve for each sample and replicates.

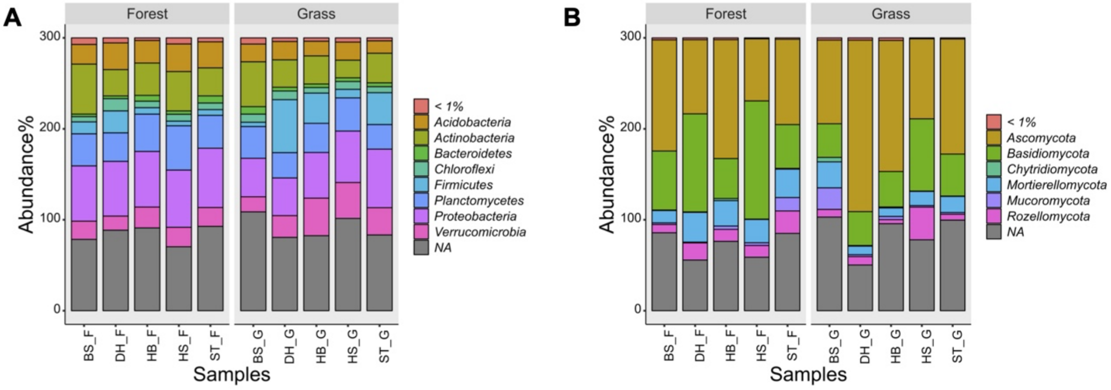


**Fig. S3** (A) Bacterial and (B) Fungal Shannon alpha diversity in each sampling sites.

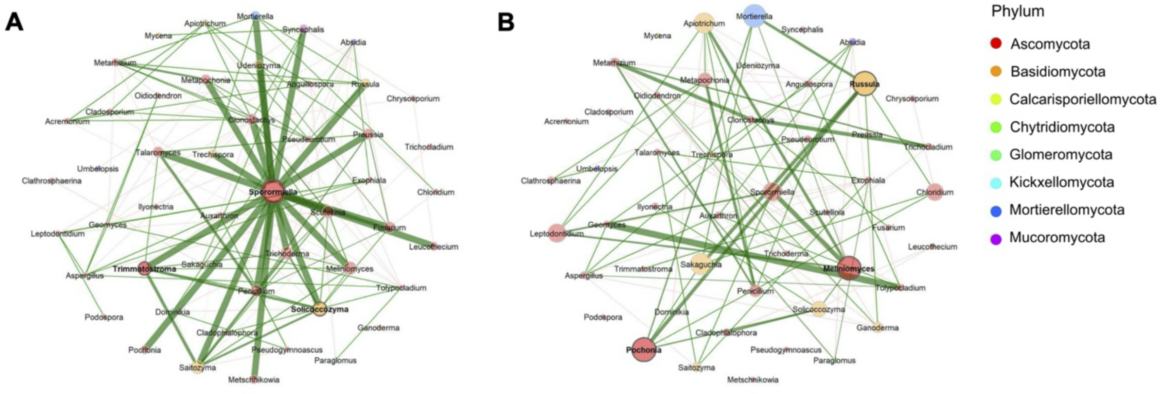


C

| | Modularity | Clustering coefficient | Positive edge % | Edge density | Natural connectivity |
|---|---|---|---|---|---|
| Grass land | 0.32 | 0.51 | 55.22 | 0.01 | 0.07 |
| Forest soil | 0.51 | 0.51 | 57.14 | 0.07 | 0.03 |

**Fig. S4** Relative abundances of bacterial (A) and fungal (B) phyla in each sampling sites.

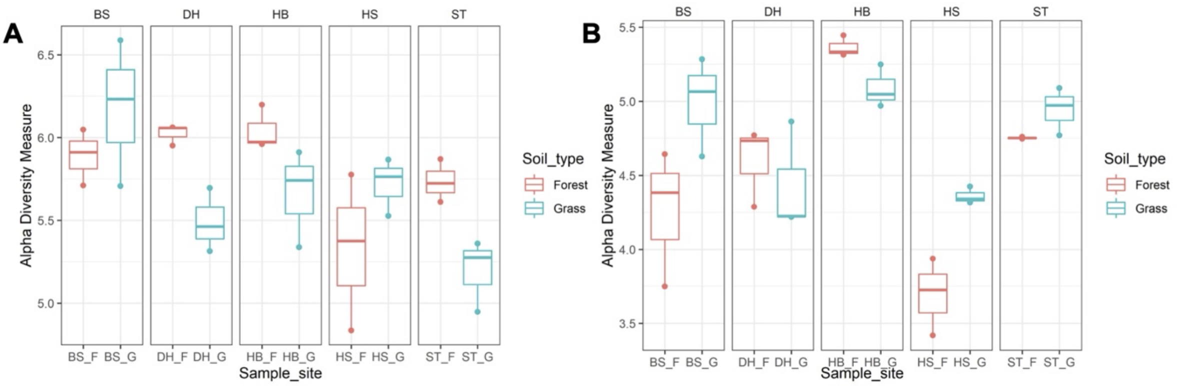


**Fig. S5** Co-occurrence comparison networks of the soil fungal community in grassland (A), and forest soil (B). Nodes represent ASVs aggregated to the genus level and are colored according to the respective phylum. ASVs that were not assigned on the genus level were annotated at a higher taxonomy level. Edges in the networks represent positive (green) and negative (red) correlations > |0.6| calculated by the SparCC method. Edges thickness is proportional to partial correlation. Eigenvector centrality of each node was used to define hub taxa and highlighted in bold. Topological characteristics of global network are listed in the table. (C) Topological characteristics of global network.
